# Struct2Graph: A graph attention network for structure based predictions of protein-protein interactions

**DOI:** 10.1101/2020.09.17.301200

**Authors:** Mayank Baranwal, Abram Magner, Jacob Saldinger, Emine S. Turali-Emre, Paolo Elvati, Shivani Kozarekar, J. Scott VanEpps, Nicholas A. Kotov, Angela Violi, Alfred O. Hero

**Author notes:** Equal contributor (Emine S. Turali-Emre, Paolo Elvati).

## Abstract

**Background:** Development of new methods for analysis of protein-protein interactions (PPIs) at molecular and nanometer scales gives insights into intracellular signaling pathways and will improve understanding of protein functions, as well as other nanoscale structures of biological and abiological origins. Recent advances in computational tools, particularly the ones involving modern deep learning algorithms, have been shown to complement experimental approaches for describing and rationalizing PPIs. However, most of the existing works on PPI predictions use protein-sequence information, and thus have difficulties in accounting for the three-dimensional organization of the protein chains.

**Results:** In this study, we address this problem and describe a PPI analysis based on a graph attention network, named *Struct2Graph*, for identifying PPIs directly from the structural data of folded protein globules. Our method is capable of predicting the PPI with an accuracy of 98.89% on the balanced set consisting of an equal number of positive and negative pairs. On the unbalanced set with the ratio of 1:10 between positive and negative pairs, Struct2Graph achieves a five-fold cross validation average accuracy of 99.42%. Moreover, Struct2Graph can potentially identify residues that likely contribute to the formation of the protein-protein complex. The identification of important residues is tested for two different interaction types: (a) Proteins with multiple ligands competing for the same binding area, (b) Dynamic protein-protein adhesion interaction. Struct2Graph identifies interacting residues with 30% sensitivity, 89% specificity, and 87% accuracy.

**Conclusions:** In this manuscript, we address the problem of prediction of PPIs using a first of its kind, 3D-structure-based graph attention network (code available at https://github.com/baranwa2/Struct2Graph). Furthermore, the novel mutual attention mechanism provides insights into likely interaction sites through its unsupervised knowledge selection process. This study demonstrates that a relatively low-dimensional feature embedding learned from graph structures of individual proteins outperforms other modern machine learning classifiers based on global protein features. In addition, through the analysis of single amino acid variations, the attention mechanism shows preference for disease-causing residue variations over benign polymorphisms, demonstrating that it is not limited to interface residues.

## Introduction

Protein–protein interactions (PPIs) are fundamental to many biological processes. Analysis of the human proteome suggests that the majority of proteins function not alone but rather as part of multi-unit complexes [1]. Indeed, PPIs are the central part of signal transduction, metabolic regulation, environmental sensing, and cellular organization [2]. In these processes, PPIs can alter enzyme kinetics, facilitate substrate channeling, form new binding sites, render a protein inactive, or modify the specificity of a protein with respect to a substrate [3]. Due to the ubiquitous presence of PPIs in living systems, being able to characterize these interactions promises to further our understanding of cellular processes [4] and provide an indispensable tool for disease treatment and drug discovery [5, 6]. PPI and their mathematical description are also essential for creation of protein analogs from other nanoscale building blocks, including but not limited to, lipids [7], sugars [8], polymers [9], nanoscale conjugates [10], and inorganic nanoparticles [11, 12, 13].

A number of strategies have been employed to decode PPIs. Traditionally, high throughput experimental techniques such as two-hybrid screens [14], tandem-affinity purification [15], and mass spectrometry [16] have been applied to create protein interaction networks. Concerns about insufficient accuracy [17], low experimental throughput [18] and high cost [19] of these methods, however, have motivated computational approaches that can complement traditional and robotic experimental protocols. Computational methods can predict whether proteins will interact based on data for the proteins’ genetic context, amino acid sequences, or structural information. Genomics analyses consider factors such as gene fusion [20], conservation across common species (phylogenetic profiling) [21], and evolutionary history [22] when determining if a pair of proteins interact.

Typical computational techniques for PPI analysis use the amino acid sequences of the two proteins to determine whether interactions occur [23, 24]. A number of features such as frequency of common sub-sequences [25] and auto-covariance [26] have been proposed to convert sequences of different lengths into a uniformly sized representation. Sequence based methods have recently been able to leverage protein databases and machine learning techniques to make high accuracy predictions. Three-dimensional (3D) structure of protein-protein complexes from sequence can be predicted by CO-threading algorithm, (COTH) that recognizing templates of protein complexes from solved complex structural databases. COTH aligns amino acid chain sequences using scoring functions and structural information [27]. The DeepPPI model [28] predicts interactions using an artificial neural network, which takes as input a feature vector that captures the composition, distribution, and order of the sequence. DeepFE [29] uses natural language processing algorithms on amino acid sequences to create low dimensional embeddings of the sequence suitable as inputs for neural network analysis. DeepFE, in particular, has been shown to be quite effective, and achieves prediction accuracy of 94.78% and 98.77% on *S. cerevisiae* and human datasets, respectively. In fact, most deep learning based methods have been shown to achieve high PPI prediction accuracy [30, 31] owing to their significantly larger representation power. In addition to relying purely on sequence-based information, modern machine learning methods often incorporate network-level information for PPI prediction. In a PPI network, each node represents a protein, while edges between them represent interactions. Thus, predicting interactions between any two nodes is a link-prediction problem in disguise. Recent methods have leveraged the network structure, along with using vectorized representation of amino acid sequences, to obtain stronger prediction performance [32, 33, 34, 35, 36, 13].

Despite their success, the above sequence based approaches do not generalize to broader classes of chemical compounds of similar scale as proteins that are equally capable of forming complexes with proteins that are not based on amino acids, and thus lack of an equivalent sequence-based representation. While the interaction of proteins with DNA can be accurately predicted [37], the methods for machine learning-based predictions for protein complexes with high molecular weight lipids [7], sugars [8], polymers [9], dendrimers [38] and inorganic nanoparticles [11, 12] that receive much attention in nanomedicine and nanodiagnostics [39, 40], are not widely known among experimentalists [41, 42, 43, 44, 45, 46, 47], although substantial strides in this direction were made with the development of unified structural descriptors for proteins and nanoparticles [13]. As a consequence, predictive computational approaches that take into account the structure of proteins and their variable non-proteinatious, biomimetic, and non-biological counterparts become possible. Some methods predict interactions using the 3D structure of the proteins [48, 49] use a knowledge-based approach to assess the structural similarity of candidate proteins to a template protein complex. As this methodology requires detailed information on the larger complex, template-free docking approaches [50] analyze the unbound protein components and identify the most promising interactions from a large set of potential interaction sites. While docking methods have shown success for some proteins, they face difficulty with proteins undergoing conformational changes during interaction [51]. Many of these structural approaches have also served as the basis for machine learning models. Zhang et al. developed PrePPI [52] which uses amino acid sequence, and phylogenetic features as inputs to a naive Bayes classifier. Northey et al. developed IntPred [53] which segments proteins into a group of patches that incorporates 3D structural information into a feature set to predict interaction with a multi-layer perception network. These models are trained on carefully curated interaction databases describing both binary interactions between proteins, and corresponding interfacing sites or atoms.

In this work, we make the first step toward a generalized method to assess supramolecular interactions of proteins with other nanostructures. The proposed method determines the probability of formation of protein-protein complexes on a nanoscale representation of proteins from crystallographic data, as contrasted to amino-acid amino sequence information. We develop a mutual graph attention network and a corresponding computational tool, *Struct2Graph*, to predict PPIs solely from 3D structural information. Instead of using several protein specific features, such as, hydrophobicity, solvent accessible surface area (SASA), charge, frequency of *ngrams*, etc., Struct2Graph uses a graph based representation of a protein globule obtained using *only* the 3D positions of atoms. This graph based interpretation allows for neural message passing [54] for efficient representation learning of proteins. Struct2Graph builds upon our prior work on metabolic pathway prediction [55], where it is shown that an equivalent graph-based structural representation of small molecules and peptides coupled with graph convolutional network, significantly outperforms other classifiers that involve computing various biochemical features as inputs. This approach also leverages generalization of graph theory to describe complex nanoscale assemblies similar to PPI [56].

Beyond the high accuracy of its PPI predictions, Struct2Graph offers a number of advantages. Similarly to the ML algorithms exploiting the idea of geometrical biomimetics, Struct2Graph only requires the 3D structure of individual proteins. Furthermore, while in this paper we focus on protein interactions, by using only the positions of atoms in our analysis, this framework can be generalized to other molecular structures where 3D information is available. Moreover, Struct2Graph is also able to provide insight into the nature of the protein interactions. Through its attention mechanism, the model can potentially identify residues that likely contribute to the formation of the protein-protein complex. Unlike other models, Struct2Graph is able to produce this data in an unsupervised manner and thus does not require protein complex information which are often unavailable [57].

The key contributions of the proposed work can be summarized as:

- **Graph convolutional network for PPI prediction:** Struct2Graph uses a multi-layer graph convolutional network (GCN) for PPI prediction from the structural data of folded protein globules. The proposed approach is general and can be applied to other nanoscale structures where 3D information is available.
- **Curation of PPI database:** A large PPI database comprising of only direct/physical interaction of *non-homologous* protein pairs^[1]^ is curated, along with information on the corresponding PDB files. Special emphasis is based on curation of PDB files based on the length of the chain ID and highest resolution within each PDB file to ensure capturing of the most complete structure information of the protein of interest.
- **State-of-the-art prediction performance:** Our method is capable of correctly predicting the PPIs with an accuracy of 98.89% on the balanced set consisting of an equal number of positive and negative pairs. On the unbalanced set with the ratio of 1:10 between positive and negative pairs, Struct2Graph achieves a five-fold cross validation average accuracy of 99.42%. Struct2Graph outperforms not only the classical feature-based machine learning approaches, but also other modern deep-learning approaches, such as Deep-PPI and DeepFE-PPI that use sequence information and feature selection for PPI prediction.
- **Unsupervised prediction of important residues:** The novel mutual attention mechanism can potentially identify important residues for the formation of the protein-protein complex. This importance can stem from either direct participation in the interaction process (i.e., binding site) or indirectly through contribution to appropriate protein folding that allows formation of the correct binding site geometry. The identification of important residues is tested for two different interaction types (neither part of the training set): (a) Proteins with multiple ligands competing for the same binding area, (b) Dynamic protein-protein adhesion interaction. Struct2Graph identifies interacting residues with 30% sensitivity, 89% specificity, and 87% accuracy.
- **Analysis of single amino acid variation (SAV) dataset:** Disease-causing mutations are known to be preferentially located within the interface core, as opposed to the rim. Of the known 2724 disease-causing SAVs and 1364 polymorphisms, our attention mechanism identifies 33.55% of all disease-causing SAVs as important (attention weights within top-20%), while 85.30% of all polymorphisms are identified as *un*important by the proposed attention mechanism, indicating significant overlap between the previously established SAV study and the important residues identified by the proposed attention mechanism.

## Materials and methods

### PPI Database

Struct2Graph focuses on structure-based predictions and interaction sites of the protein pairs. Our PPI database is therefore produced based on only direct/physical interactions of proteins. To build a large physical interaction database, comprising of only *heterologous* pairs, we searched all possible databases available (STRING, BioGRID, IntAct, MINT, BIND, DIP, HPRD, APID, OpenWetWare). Not all PPI databases use the same publications and same ontologies to report the interactions. Consequently, it is not surprising that each database reports PPI differently. Therefore, only up to a 75% concordance between all PPI databases is achieved [58]. For Struct2Graph, two of the largest compiled databases, IntAct [59] and STRING [60] are chosen for further analysis, and results are compared to each other to find the true interactions. Only concordant matches between these two databases are chosen. Struct2Graph database is compiled from commonly studied organism’s (*Saccharomyces cerevisiae, Homo sapiens, Escherichia coli, Caenorhabditis elegans* and *Staphylococcus aureus*) PPIs. For these organisms, IntAct provides 427,503 PPIs, and STRING provides 852,327 PPIs.

STRING distinguishes the type of interactions as “activation”, “binding”, “catalysis”, “expression”, “inhibition” and “reaction”. IntAct, on the other hand, describe the type of interactions as “association”, “physical association”, “direct association/interaction”, and “colocalization”. Only “direct association/interactions” from IntAct and “binding” from STRING were considered as physical interactions. We only choose concordant pairs of physical interactions from both databases. Therefore, extracting only physical interaction data from the rest of the interactions reduces the actual number of PPIs to 12,676 pairs for IntAct and 446,548 pairs for STRING. Negative PPI is extracted from the work that derives negative interaction from large-scale two-hybrid experiments [61]. The negative protein-protein pairs from the two-hybrid system are compared further with the database constructed from STRING and INTact, and only the pairs that are not involved in any interaction at all, are chosen. We further exclude the co-localized protein pairs in our analysis. Structure information for Struct2Graph is obtained from PDB files. Hence, we only used the pairs which have associated PDB files. This reduces the total number of pairs to 117,933 pairs (4698 positive and 112,353 negatives). Some proteins are well-studied as they are in the scope of current medical and biotechnological interest. As a result, there is more than one cross-reference to PDB files since various structures are accessible for these proteins. To find the proteins matched with PDB files, all proteins from the database are matched with UniProt accession numbers (UniProt Acc) and mapped with PDB files in UniProt [62]. Unfortunately, not all proteins are crystallized fully in each PDB file, and random choice of PDB file may cause incomplete information of the binding site of the protein. Therefore, we curated the PDB files based on the length of the chain ID and highest resolution within each PDB file to ensure that we capture the most complete structure information of the protein of interest. The chain length and the resolution of each protein’s crystal structure were obtained from the RCSB website [63]. The complete set of negative pairs was reduced to 5036 pairs to create a fairly balanced training sample with an approximately equal number of positive and negative pairs. For this curated database consisting of only heterologous pairs, we defined two classes, “0” for non-interacting (negative: not forming complexes) pairs and “1” for interacting (positive: forming complexes) pairs.

### Mutual graph attention network for protein-protein pairs

We present a novel multi-layer mutual graph attention network (GAT) based architecture for PPI prediction task, summarized in Fig 1. We refer to this architecture as *Struct2Graph*, since the inputs to the proposed GAT are coarse grained structural descriptors of a query protein-protein pair. Struct2Graph outputs the probability of interaction between the query proteins. Struct2Graph uses two graph convolutional networks (GCNs) with weight sharing, and a mutual attention network to extract relevant geometric features related to query protein pairs. These extracted features are then concatenated and fed to a feedforward neural network (FNN) coupled with a `SoftMax` function, which finally outputs a probability of the two classes - ‘0’ (negative pairs) and ‘1’ (positive pairs). This section first describes the preprocessing and fingerprinting procedure specifying how spatial information on protein pairs are converted into corresponding protein graphs, and then elaborates on different components of the Struct2Graph deep learning architecture.

**Figure 1.**
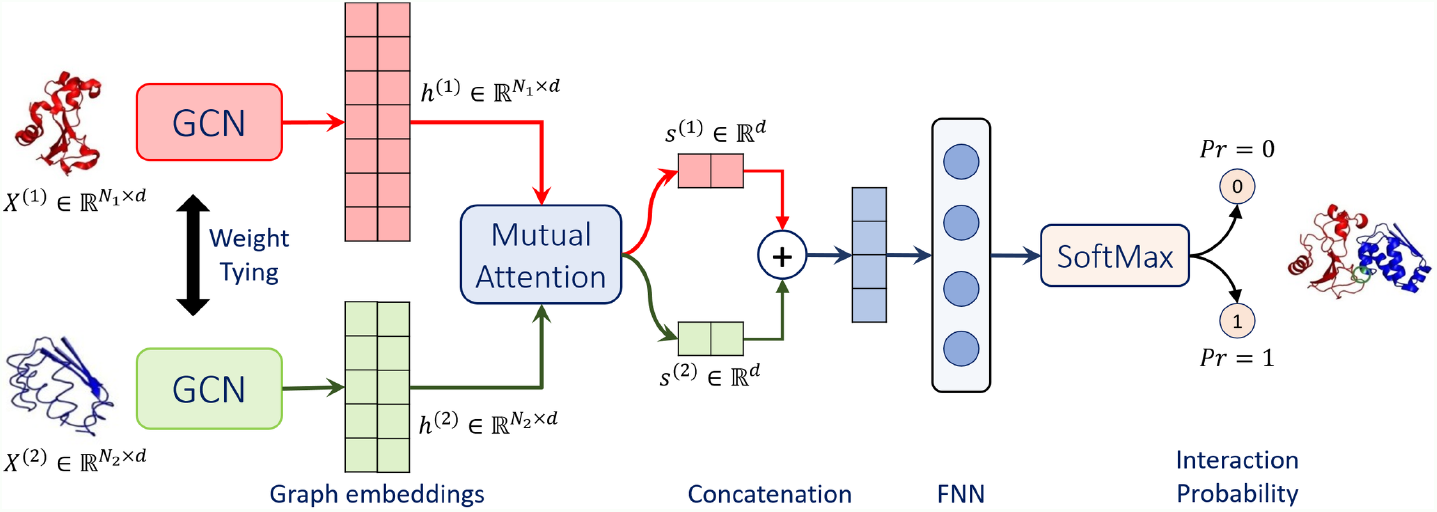
Struct2Graph schematic. Struct2Graph graph convolutional network (GCN) for incorporating mutual attention for PPI prediction. The GCN classifies whether or not a protein pair (*X*^(1)^ and *X*^(2)^ on far left) interacts and predicts the interaction sites (on far right).

#### Protein structure graph

The purpose of the graph construction step is to capture the salient geometry of the proteins in a way that is amenable to further dimensionality reduction by the neural network. There are many possible ways of constructing a graph from spatial coordinates of individual atoms, and each captures a different level of detail about the geometry of the protein [13]. For instance, the protein-contact graph described in [64] adds edges between three nearest neighbors and identifies the nodes as helices, sheets, and turns. Ralaivola et al. [65] uses molecular fingerprints to prescribe contact graphs for chemical compounds. Pires et al. [66] employs a distance-based approach for constructing protein graphs by encoding distance patterns between constituent atoms. Cha et al. [13] pioneered multidimentional protein graphs with embedded chemical, geometrical and graph theoretical descriptors. Our approach to constructing protein graphs is inspired by the latter, however, it is generalizable to other non-protein structures as well. We first aggregate atoms into the amino acids that they constitute and define the position of an amino acid to be the average of the positions of its constituent atoms. These amino acids form the vertices of the protein graph. An edge is placed between two vertices if the distance between them is less than some threshold. Unlike the previous studies with 7Å threshold [13], in this work, we use a threshold of 9.5Å for creating a protein graph from the mean positions of amino acids. This threshold was obtained empirically so as to render the underlying graph fully connected, while simplifying the graph representation. Note that while we use amino acids as constituent vertices of the protein graphs, the approach can be easily extended to multiresolution representation, where a vertex represents two or more amino acids. The coarse-grained representation opens up new possibilities for studying other nanoscale components of protein complexes, such as, lipids and polysaccharides, since, lowering the level of representation from all-atom to submolecular can be easily generalized to other non-protein entities. Graphs with greater structural refinement can also be obtained by using functional groups as amino acids. Moreover, this geometric construction of protein graphs ensures that salient geometric features, such as spatial proximity of non-adjacent amino acids along the polypeptide chain are captured. A sequence based representation of proteins might not capture this geometrical structure as well (see Fig 2).

**Figure 2.**
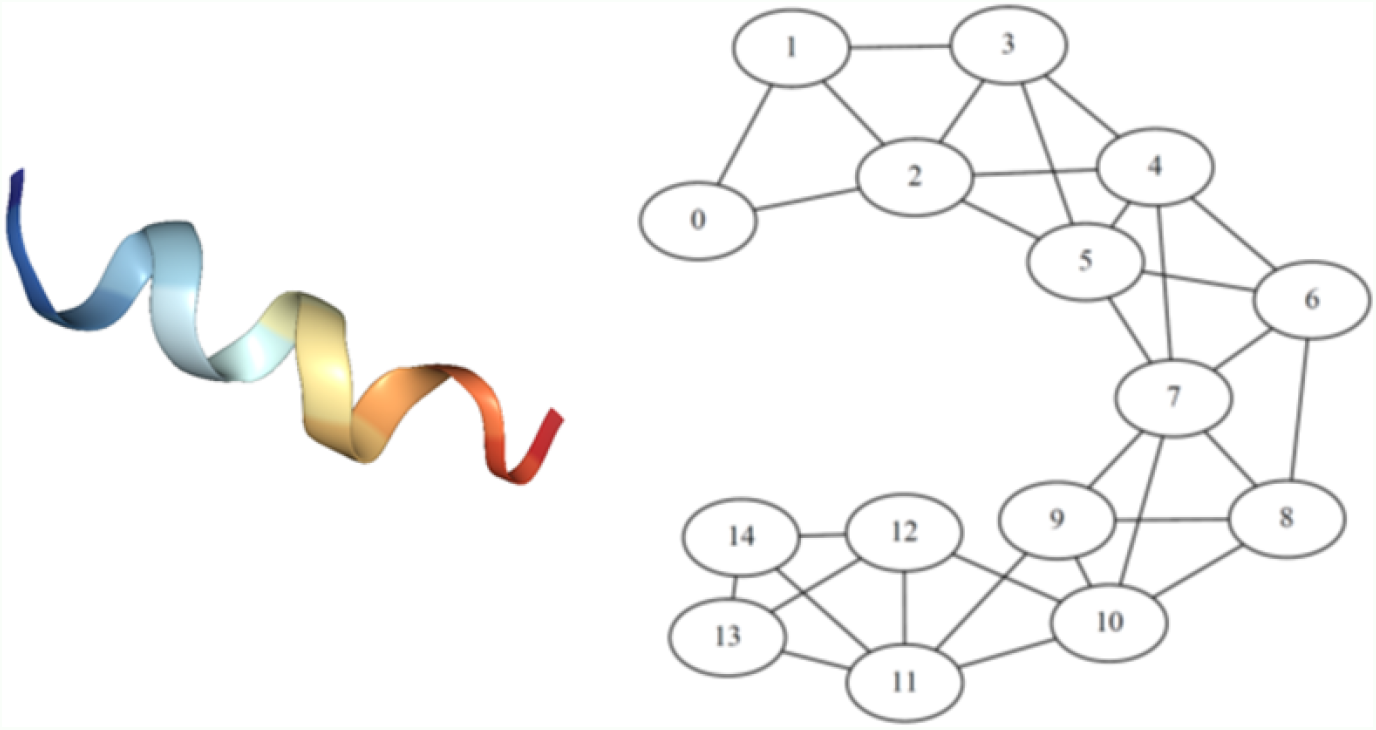
Protein and protein graph. Illustration of extracted protein structure graph (right) from the corresponding PDB description of a peptide segment (left) of the *S. cerevisiae* alpha-factor receptor. The graph is extracted by thresholding the distances between amino acids. The helical structure of the protein (left) gets captured in the corresponding protein graph (right) where, for example, amino acid 4 is linked with amino acid 7.

The graph construction approach converts spatial information associated with a protein into an equivalent protein graph object 𝒢 = (𝒱, ℰ), where 𝒱 is the set of vertices and ℰ is the set of edges between them. In the context of protein graph in Fig 2, *v*_*i*_ ∈ 𝒱 is the *i*^th^ amino acid and *e*_*ij*_ ∈ ℰ represents an edge between *i*^th^ and *j*^th^ amino acids, satisfying their proximity within the specified threshold of 9.5Å. These graph objects must be embedded into real-valued vector space in order to employ our machine learning framework. We use 1-neighborhood subgraphs [55] induced by the neighboring vertices and edges at 1-hop distance from a vertex. A dictionary of all unique subgraphs is constructed by scanning all protein graphs in the training database. Thus, each vertex within a protein is equivalently represented by an element in the dictionary.

#### Graph convolutional network acting on protein graphs

A graph convolutional network (GCN) maps graphs to real-valued *embedding vectors* in such a way that the geometry of the embedding vectors reflects similarities between the graphs. The embedding portion of the GCN works as follows. To each vertex *v*_*i*_ ∈ 𝒱, we associate a *d*-dimensional feature vector, which encodes the 1-neighborhood subgraph induced by the neighboring vertices and edges at 1-hop distance from a vertex. This is in contrast to explicit inclusion of amino acid specific features, such as, hydrophobicity, solvent accessible surface area (SASA), charge, etc. In our encoding, similar to other studies [67, 55], each element of the dictionary of subgraphs is assigned a random unit-norm vector.

Each layer of the GCN updates all vertex features by first replacing each vertex feature by a normalized average over vertex features of all 1-hop neighboring vertices. This is followed by an affine transformation given by the trained weight matrices and bias parameters. In order to impart expressivity to the GCN architecture, each coordinate of the resulting affine transformed embedding vector is passed through a nonlinear activation function, such as, rectified linear unit (ReLU) or sigmoid activations. This process is repeated for all the subsequent layers, and the output of the final layer is the newly transformed embedding (feature) vector that is propagated further to the mutual attention network. Here, the number of layers is a hyperparameter, while the weight matrices are learned from the training data in order to optimize performance of the entire system on the interaction prediction task.

More concisely, given input protein graphs 𝒢^(1)^, 𝒢^(2)^ with adjacency matrices *A*^(1)^, *A*^(2)^ consisting of *N*_1_, *N*_2_ vertices (amino acids), and quantities 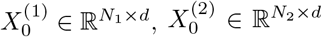 representing the *d*-dimensional embedding of the vertex subgraphs of the query protein-protein pair, respectively, an *l*-layer GCN updates vertex embeddings using the following update rule:

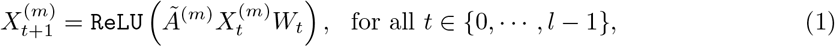

where 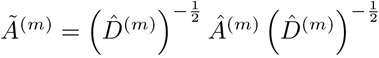 denotes the normalized adjacency matrices, and *m* ∈ *{*1, 2*}*. Here, *Â*^(*m*)^ = *A*^(*m*)^ + *I* and 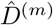 is the degree matrix of *Â*^(*m*)^. Parameters *W*_*t*_ denote the weight matrix associated with the *t*^th^-layer of the GCN. The feature embeddings 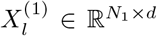 and 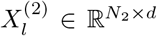 produced by the final layer of GCN are fed to a mutual attention network and hereafter denoted as *h*^(1)^ and *h*^(2)^, respectively, for notational convenience.

#### Mutual attention network for PPI prediction

The purpose of the proposed mutual attention network is two fold: (a) extract relevant features for the query protein-protein pair that *mutually* contribute towards prediction of physical interaction of proteins, (b) combine embedding matrices of dissimilar dimensions *N*_1_ *× d* and *N*_2_ *× d* to produce a representative single output embedding vector of dimension (2*d*). Attention mechanisms were originally introduced for interpreting sequence-to-sequence translation models by allowing the models to attend differently to different parts of the encoded inputs. Since then, it has been adapted in other fields of deep learning, such as, computer vision [68], and bioinformatics [67].

The mutual attention mechanism proposed in this work computes attention weights 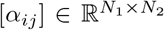 and context vectors *s*^(1)^ ∈ ℝ^*d*^, *s*^(2)^ ∈ ℝ^*d*^ from the GCN-transformed hidden embeddings *h*^(1)^ and *h*^(2)^. The attention weights are computed as:

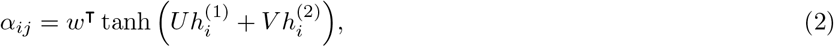

where *U, V* ∈ ℝ ^*d×d*^ and *w* ∈ ℝ^*d*^ are parameters of the mutual attention network that are trained in an end-to-end fashion along with the weights of the GCN. These attention weights are then translated to context vectors *s*^(1)^, *s*^(2)^ using the following knowledge selection procedure:

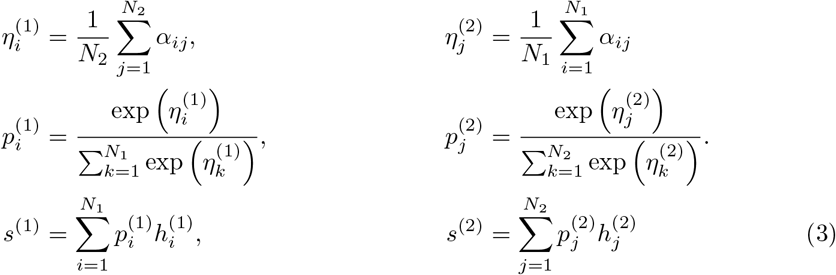

Here 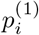 and 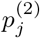^)^ denote the relative weights of the amino acids *i* and *j* of the query protein-protein pair that contribute towards interaction prediction. Those vertices whose learned attention weights are large are likely to represent residues that participate directly or indirectly towards forming protein-protein complex.

The context vectors *s*^(1)^ and *s*^(2)^ are then concatenated into a single context vector of dimensions 2*d*, which is used as input to a single-layer, fully connected feedforward neural network (FNN) represented by *f* (*·*) to produce a two-dimensional output vector. The FNN is parameterized by another weight matrix to be learned in an end-to-end manner. A final SoftMax layer is applied to produce a probability, one for each of the possible classes: 0 or 1, as shown in Equation (4). This output represents the classifier’s prediction of the probability that the two input proteins interact.

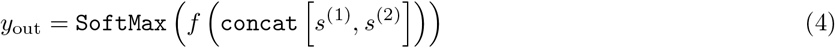

## Results

As part of our assessment, we compare the performance of Struct2Graph for PPI predictions against a number of recent machine learning models. These methods include: (a) DeepFE model [29], where we train the natural language processing network on the same database used in the original publication and feed the embeddings into a fully connected feedforward neural network. (b) DeepPPI [28], where we extract 1164 sequence features related to the amino acid composition, distribution, and order. A separate neural network is used for each protein in the protein-protein pair and their outputs are concatenated into a final network for classification. Furthermore, as was done in the original publication [28], we implement these features into a number of traditional machine learning models [69], such as (c) Gaussian naive Bayes (GaussianNB) classifier, (d) Quadratic discriminant analysis (QDA), (e) *k*-nearest neighbor (*k*-NN) classifier, (f) Decision tree (DT) classifier, (g) Random forest (RF) classifier, (h) Adaboost classifier, and (i) Support vector classifier (SVC) [70]. All models are implemented in Python 3.6.5 on an Intel i7-7700HQ CPU with 2.8GHz x64-based processor. For common machine learning classifiers, such as, GaussianNB, QDA, SVC, RF, DT, *k*-NN and Adaboost, we use the readily available implementation in the scikit-learn [69] module. Deep learning classifiers, in particular, DeepPPI [71] and DeepFE-PPI [72] are implemented in Keras [73], while Struct2Graph is implemented in PyTorch [74].

For Struct2Graph, the hyperparameters of the models are tuned in order to achieve the reported accuracies. The tuning is obtained by performing grid search over the set of possible hyperparameter settings. The hyperparameters of our Struct2Graph implementation are as follows: `optimizer`: Adam optimizer [75] with learning rate *λ* = 10^*−*3^ and rate-decay of 0.5 per 10 epochs; `total epochs`: 50; `number of GCN layers`: *l* = 2; `GCN embedding dimension`: *d* = 20; `loss function`: binary cross-entropy. For other competing methods, we use the tuned hyperparameters that are adopted from the original publications.

### Performance on balanced databases

Table 1 summarizes the comparisons of Struct2Graph and various machine learning models for PPI prediction for a five-fold stratified cross validation study. In the cross validation, the 10004 pairs (4698 positive and 5036 negatives) are randomly partitioned into five subsamples of equal size. Of these five subsamples, a single subsample is retained as the validation data for testing various machine learning models, and the remaining four subsamples are used as training data. In order to reduce the training time with our Struct2Graph model, 800 pairs are randomly sampled with replacement among the 8003 pairs (80%) in each epoch, and the performance on the randomly chosen 800 pairs is used to update the parameters of the neural network. This modification not only reduces the training time considerably, but also injects noise into the training data to avoid any potential overfitting.

**Table 1.** Five-fold cross-validation performance analysis of several machine learning methods on balanced dataset (1:1). Note that the proposed Struct2Graph method outperforms all other methods on the majority of metrics.

| Method | Performance (%) on Balanced training set - 1:1 |  |  |  |
| --- | --- | --- | --- | --- |
|  | Accuracy | Precision | Recall | Specificity |
| GaussianNB | 72.14±2.91 | 98.41±0.51 | 45.05±6.10 | 99.24±0.30 |
| QDA | 78.66±3.44 | 70.43±3.41 | **99.42±0.40** | 57.90±7.12 |
| *k*-NN | 94.19±0.56 | 99.49±0.08 | 88.83±1.10 | 99.54±0.07 |
| Decision Trees | 96.20±0.43 | 97.59±0.28 | 94.75±0.99 | 97.66±0.29 |
| Random Forest | 98.86±0.29 | 99.45±0.19 | 98.27±0.49 | 99.45±0.19 |
| Adaboost | 97.85±0.26 | 98.76±0.18 | 96.92±0.51 | 98.78±0.18 |
| SVC | 98.49±0.33 | 99.44±0.18 | 97.53±0.61 | 99.45±0.18 |
| DeepPPI | 97.22±0.44 | 98.26±0.82 | 96.14±0.88 | 98.29±0.83 |
| DeepFE-PPI | 98.64±0.32 | 99.16±0.28 | 98.12±0.51 | 99.17±0.28 |
| Struct2Graph | **98.89±0.24** | **99.50±0.36** | 98.37±0.34 | **99.45±0.42** |

| Method | MCC | F1-score | ROC-AUC | NPV |
| --- | --- | --- | --- | --- |
| GaussianNB | 52.69±4.38 | 61.53±6.00 | 95.24±0.33 | 64.46 ± 2.37 |
| QDA | 63.06±5.23 | 82.40±2.27 | 78.66±3.43 | **99.05±0.58** |
| *k*-NN | 88.89±1.02 | 93.86±0.63 | 98.79±0.21 | 89.92±0.89 |
| Decision Trees | 92.45±0.84 | 96.15±0.46 | 96.36±0.30 | 95.23±0.61 |
| Random Forest | 97.74±0.58 | 98.86±0.30 | **99.63±0.08** | 98.30±0.46 |
| Adaboost | 95.72±0.52 | 97.83±0.27 | 99.20±0.08 | 96.94±0.50 |
| SVC | 97.01±0.66 | 98.48±0.34 | **99.63±0.09** | 97.58±0.59 |
| DeepPPI | 94.47±0.87 | 97.19±0.44 | 99.28±0.11 | 96.23±0.81 |
| DeepFE-PPI | 97.29±0.64 | 98.64±0.32 | 99.52±0.09 | 98.14±0.50 |
| Struct2Graph | **97.79±0.49** | **98.94±0.20** | 99.55±0.16 | 98.24±0.42 |

The performance is reported for various measures, such as, accuracy, precision, recall, specificity or the true negative rate, Matthews correlation coefficient (MCC), F_1_-score, area under the receiver operating characteristic curve (ROC-AUC), and negative predictive value (NPV) (see Tables 1-10). For a balanced training set (Table 1), Struct2Graph outperforms any other existing machine learning models in the literature for all the measures (except for the recall, NPV, and ROC-AUC scores) with an average accuracy and precision of 98.89% and 99.50%, respectively. This is despite the fact that we significantly downsample the number of pairs in each epoch during the training process of the proposed Struct2Graph model.

**Table 2.** Bootstrap resampling performance analysis of several machine learning methods on balanced dataset (1:1). Note that the proposed Struct2Graph method outperforms all other methods on the majority of metrics.

| Method | Performance (%) on Balanced training set - 1:1 |  |  |  |
| --- | --- | --- | --- | --- |
|  | Accuracy | Precision | Recall | Specificity |
| GaussianNB | 71.90±3.19 | 99.01±0.38 | 46.39±5.95 | 99.47±0.23 |
| QDA | 54.46±0.93 | 53.28±0.77 | **99.96±0.06** | 5.30±1.74 |
| k-NN | 92.77±0.22 | 99.38±0.13 | 86.62±0.50 | 99.41±0.11 |
| Decision Trees | 95.15±0.51 | 97.13±0.38 | 93.41±0.88 | 97.03±0.31 |
| Random Forest | 98.70±0.11 | 99.39±0.18 | 98.10±0.14 | 99.35±0.18 |
| Adaboost | 96.61±0.25 | 97.83±0.40 | 95.60±0.21 | 97.71±0.44 |
| SVC | 98.05±0.19 | 99.35±0.15 | 96.87±0.28 | 99.32±0.15 |
| DeepPPI | 97.16±0.40 | 98.04±0.96 | 96.27±0.84 | 98.06±0.98 |
| DeepFE-PPI | 98.41±0.12 | 98.81±0.50 | 98.11±0.45 | 98.73±0.55 |
| Struct2Graph | **98.96±0.19** | **99.40±0.09** | **98.57±0.35** | **99.47±0.09** |
| Method | MCC | F1-score | ROC-AUC | NPV |
| GaussianNB | 53.40±4.44 | 62.93±5.81 | 96.04±0.12 | 63.30 ± 2.72 |
| QDA | 16.43±2.53 | 69.50±0.66 | 52.63±0.84 | **99.37±0.83** |
| k-NN | 86.36±0.40 | 92.56±0.30 | 98.32±0.16 | 87.31±0.38 |
| Decision Trees | 95.23±0.55 | 95.27±0.54 | 94.21±0.71 | 93.18±0.81 |
| Random Forest | 97.41±0.22 | 98.74±0.11 | 99.58±0.07 | 97.97±0.17 |
| Adaboost | 93.25±0.50 | 96.70±0.23 | 99.00±0.09 | 95.36±0.29 |
| SVC | 96.13±0.37 | 98.10±0.20 | 99.53±0.09 | 96.71±0.28 |
| DeepPPI | 94.36±0.81 | 97.14±0.40 | 99.05±0.14 | 96.34±0.78 |
| DeepFE-PPI | 96.82±0.25 | 98.45±0.12 | 99.52±0.05 | 97.99±0.20 |
| Struct2Graph | **97.91±0.38** | **98.98±0.19** | **99.62±0.17** | 98.50±0.33 |

**Table 3.** Five-fold cross-validation performance analysis of deep-learning based machine learning methods on unbalanced dataset (1:2).

| Method | Performance (%) on Unbalanced training set - 1:2 |  |  |  |
| --- | --- | --- | --- | --- |
|  | Accuracy | Precision | Recall | Specificity |
| DeepPPI | 97.40±0.44 | 98.64±0.61 | 93.52±1.64 | 99.35±0.30 |
| DeepFE-PPI | 98.91±0.09 | 99.00±0.32 | 97.71±0.33 | 99.51±0.16 |
| Struct2Graph | **99.03±0.24** | **99.13±0.25** | **98.11±0.58** | **99.53±0.13** |
| Method | MCC | F1-score | ROC-AUC | NPV |
| DeepPPI | 94.16±0.97 | 96.00±0.72 | 99.19±0.21 | 96.85±0.76 |
| DeepFE-PPI | 97.54±0.20 | 98.35±0.13 | **99.56±0.08** | **99.86±0.16** |
| Struct2Graph | **97.87±0.51** | **98.62±0.32** | 99.47±0.20 | 98.97±0.34 |

**Table 4.** Five-fold cross-validation performance analysis of deep-learning based machine learning methods on unbalanced dataset (1:3).

| Method | Performance (%) on Unbalanced training set - 1:3 |  |  |  |
| --- | --- | --- | --- | --- |
|  | Accuracy | Precision | Recall | Specificity |
| DeepPPI | 98.19±0.58 | 98.73±0.40 | 93.98±2.43 | 99.59±0.13 |
| DeepFE-PPI | 98.96±0.27 | 98.30±0.46 | 97.52±0.88 | 99.44±0.15 |
| Struct2Graph | **99.30±0.22** | **99.17±0.44** | **98.19±1.09** | **99.71±0.13** |
| Method | MCC | F1-score | ROC-AUC | NPV |
| DeepPPI | 95.15 ± 1.55 | 96.28 ± 1.24 | 99.27±0.14 | 98.03±0.78 |
| DeepFE-PPI | 97.21±0.72 | 97.90±0.55 | **99.51±0.11** | 99.18±0.29 |
| Struct2Graph | **98.20±0.56** | **98.67±0.41** | 99.49±0.21 | **99.33±0.38** |

**Table 5.** Five-fold cross-validation performance analysis of deep-learning based machine learning methods on unbalanced dataset (1:5).

| Method | Performance (%) on Unbalanced training set - 1:5 |  |  |  |
| --- | --- | --- | --- | --- |
|  | Accuracy | Precision | Recall | Specificity |
| DeepPPI | 97.78±0.45 | 98.33±0.39 | 88.20±2.74 | **99.70±0.07** |
| DeepFE-PPI | 98.97±0.27 | 98.19±0.49 | 95.60±1.52 | 99.65±0.10 |
| Struct2Graph | **99.13±0.18** | **98.49±0.85** | **96.63±0.93** | 99.68±0.19 |
| Method | MCC | F1-score | ROC-AUC | NPV |
| DeepPPI | 91.87±1.68 | 92.97±1.53 | 98.69±0.50 | 97.69±0.52 |
| DeepFE-PPI | 96.28±1.00 | 96.87±0.85 | **99.56±0.25** | 99.12±0.30 |
| Struct2Graph | **97.03±0.56** | **97.55±0.45** | 99.17±0.25 | **99.26±0.23** |

**Table 6.** Five-fold cross-validation performance analysis of deep-learning based machine learning methods on unbalanced dataset (1:10).

| Method | Performance (%) on Unbalanced training set - 1:10 |  |  |  |
| --- | --- | --- | --- | --- |
|  | Accuracy | Precision | Recall | Specificity |
| DeepPPI | 98.24±0.49 | 95.83±2.60 | 84.33±4.23 | 99.63±0.23 |
| DeepFE-PPI | 99.17±0.33 | 96.56±1.09 | 94.19±2.87 | 99.67±0.10 |
| Struct2Graph | **99.42±0.14** | **97.54±1.28** | **96.43±2.49** | **99.73±0.16** |
| Method | MCC | F1-score | ROC-AUC | NPV |
| DeepPPI | 88.95±3.14 | 89.66±2.94 | 97.18±1.24 | 98.45±0.41 |
| DeepFE-PPI | 94.91±2.07 | 95.35±1.90 | **99.48±0.32** | 99.42±0.29 |
| Struct2Graph | **96.65±1.12** | **96.96±1.07** | 99.45±0.70 | **99.63±0.22** |

**Table 7.** Bootstrap resampling performance analysis of deep-learning based machine learning methods on unbalanced dataset (1:2).

| Method | Performance (%) on Unbalanced training set - 1:2 |  |  |  |
| --- | --- | --- | --- | --- |
|  | Accuracy | Precision | Recall | Specificity |
| Decision Trees | 94.86±0.58 | 95.67±1.29 | 89.48±0.79 | 97.79±0.67 |
| Random Forest | 98.74±0.15 | 99.10±0.17 | 97.33±0.44 | 99.45±0.09 |
| DeepPPI | 97.91±0.38 | 98.37±0.69 | 95.32±1.40 | 99.21±0.34 |
| DeepFE-PPI | 98.53±0.20 | 98.41±0.62 | 97.37±0.31 | 99.16±0.31 |
| Struct2Graph | **98.91±0.24** | **99.17±0.15** | **97.89±0.17** | **99.52±0.27** |
| Method | MCC | F1-score | ROC-AUC | NPV |
| Decision Trees | 88.69±1.28 | 92.47±0.82 | 93.66±0.58 | 94.47±0.50 |
| Random Forest | 97.25±0.33 | 98.20±0.22 | 99.71±0.08 | 98.56±0.24 |
| DeepPPI | 95.29±0.85 | 96.81±0.60 | 99.29±0.25 | 97.70±0.67 |
| DeepFE-PPI | 96.76±0.45 | 97.88±0.31 | 99.41±0.17 | 98.59±0.09 |
| Struct2Graph | **97.59±0.51** | **98.43±0.30** | **99.73±0.18** | **98.87±0.16** |

**Table 8.** Bootstrap resampling performance analysis of deep-learning based machine learning methods on unbalanced dataset (1:3).

| Method | Performance (%) on Unbalanced training set - 1:3 |  |  |  |
| --- | --- | --- | --- | --- |
|  | Accuracy | Precision | Recall | Specificity |
| Decision Trees | 95.72±0.46 | 95.55±0.74 | 87.98±1.63 | 98.52±0.26 |
| Random Forest | 98.80±0.14 | 98.29±0.33 | 97.15±0.34 | 99.39±0.11 |
| DeepPPI | 97.78±0.36 | 98.43±0.42 | 92.60±1.38 | 99.51±0.13 |
| DeepFE-PPI | 98.86±0.11 | 98.42±0.54 | 97.30±0.31 | 99.43±0.20 |
| Struct2Graph | **99.01±0.16** | **98.83±0.37** | **97.42±0.51** | **99.59±0.13** |
| Method | MCC | F1-score | ROC-AUC | NPV |
| Decision Trees | 88.88±1.14 | 91.60±0.88 | 93.26±0.81 | 95.78±0.61 |
| Random Forest | 96.90±0.39 | 97.72±0.29 | 99.72±0.05 | 98.98±0.10 |
| DeepPPI | 94.04±0.98 | 95.42±0.77 | 99.06±0.47 | 97.58±0.44 |
| DeepFE-PPI | 97.08±0.28 | 97.85±0.20 | 99.34±0.11 | 99.03±0.05 |
| Struct2Graph | **97.46±0.42** | **98.12±0.32** | **99.75±0.20** | **99.08±0.18** |

**Table 9.** Bootstrap resampling performance analysis of deep-learning based machine learning methods on unbalanced dataset (1:5).

| Method | Performance (%) on Unbalanced training set - 1:5 |  |  |  |
| --- | --- | --- | --- | --- |
|  | Accuracy | Precision | Recall | Specificity |
| Decision Trees | 95.56±0.45 | 94.14±0.79 | 80.91±2.12 | 98.87±0.16 |
| Random Forest | 98.38±0.19 | 98.25±0.65 | 92.82±1.07 | 99.63±0.13 |
| DeepPPI | 97.96±0.46 | 97.13±2.71 | 90.58±3.21 | 99.44±0.57 |
| DeepFE-PPI | 98.90±0.31 | 98.20±0.29 | 95.64±1.75 | 99.61±0.07 |
| Struct2Graph | **99.16±0.17** | **98.29±0.64** | **97.08±1.01** | **99.69±0.13** |
| Method | MCC | F1-score | ROC-AUC | NPV |
| Decision Trees | 84.72±1.33 | 87.01±1.18 | 89.89±1.04 | 95.82±0.55 |
| Random Forest | 94.53±0.70 | 95.45±0.61 | 99.66±0.15 | 99.03±0.11 |
| DeepPPI | 92.58±1.66 | 93.66±1.46 | 98.96±0.13 | 98.15±0.61 |
| DeepFE-PPI | 96.24±1.11 | 96.89±0.94 | 99.49±0.11 | 99.05±0.16 |
| Struct2Graph | **97.03±0.51** | **97.53±0.41** | **99.71±0.26** | **99.40±0.23** |

**Table 10.** Bootstrap resampling performance analysis of deep-learning based machine learning methods on unbalanced dataset (1:10).

| Method | Performance (%) on Unbalanced training set - 1:10 |  |  |  |
| --- | --- | --- | --- | --- |
|  | Accuracy | Precision | Recall | Specificity |
| Decision Trees | 96.63±0.46 | 91.66±1.47 | 73.14±2.93 | 99.26±0.14 |
| Random Forest | 97.85±0.25 | 95.87±0.68 | 82.12±2.28 | 99.61±0.06 |
| DeepPPI | 98.09±0.77 | 95.30±3.86 | 83.29±7.49 | 99.58±0.37 |
| DeepFE-PPI | 98.50±0.46 | 96.56±0.38 | 87.86±5.00 | 99.66±0.05 |
| Struct2Graph | **99.26±0.15** | **97.04±0.70** | **95.59±0.73** | **99.67±0.10** |
| Method | MCC | F1-score | ROC-AUC | NPV |
| Decision Trees | 80.13±2.06 | 81.33±1.99 | 86.20±1.48 | 97.07±0.45 |
| Random Forest | 87.60±1.12 | 88.44±1.11 | 99.49±0.22 | 98.03±0.32 |
| DeepPPI | 88.01±4.96 | 88.69±4.82 | 96.65±1.60 | 98.35±0.73 |
| DeepFE-PPI | 91.27±2.80 | 91.92±2.72 | 99.50±0.20 | 98.69±0.24 |
| Struct2Graph | **95.90±0.60** | **96.31±0.52** | **99.54±0.22** | **99.50±0.12** |

Note from Table 1 that while QDA outperforms Struct2Graph in terms of recall and NPV scores, it does very poorly in terms of other measures indicating that the QDA classifier largely predicts positive interactions resulting in low false negative counts. Another observation is that the performance of Struct2Graph is only slightly better than that of another deep learning PPI model, DeepFE-PPI for this balanced training set. However, as discussed below, DeepFE-PPI does not perform as well for unbalanced training set, where positive interactions are underrepresented among all interactions, a case that often arises in practice.

The primary purpose of *k*-fold cross validation study is to measure the generalization capabilities of a model. Bootstrap resampling, on the other hand, is primarily used to establish empirical distribution functions for a widespread range of statistics. It works by performing sampling with replacement from the original dataset, and at the same time assuming that the data points that have not been chosen, are the test dataset. We repeat this procedure several times and compute the average score as estimation of the performances of various classifiers. Table 2 summarizes the comparisons of Struct2Graph and various machine learning models on a balanced dataset for PPI prediction for a bootstrap resampling method repeated over five times. As before, we downsample the number of pairs in each epoch during the training process of the Struct2Graph in order to speed up computation and avoid any potential overfitting. The performance statistics for the Struct2Graph method with bootstrap resampling are very similar to the ones obtained with a five-fold cross-validation study. Struct2Graph is shown to outperform other existing machine learning models for all the measures (except for the recall and NPV scores) with an average accuracy and precision of 98.96% and 99.40%, respectively. Interestingly, the performances of the DeepPPI and the DeepFE-PPI methods are marginally worse than that of the Random Forest classifier on the balanced set. However, as the class imbalance increases, DeepFE-PPI is shown to outperform the Random Forest classifier. We have, thus, also included the Random Forest classifier for comparative analysis on the unbalanced datasets.

### Performance on unbalanced database

In most practical scenarios, the number of negative pairs is expected to be larger than positive pairs, since only a small fraction of protein pairs interact within all possible pairs. We thus evaluate the performance of the deep learning models, Deep-PPI and DeepFE-PPI against the proposed Struct2Graph model on various unbalanced training sets, where the number of negative pairs outnumber the positive pairs. These results are summarized in Tables 3-6 for several databases with varying ratios of positive to negative pairs: (a) 1:2 (2518 positive and 5036 negative), (b) 1:3 (1679 positive and 5036 negative), (c) 1:5 (1007 positive and 5036 negative), and (d) 1:10 (504 positive and 5036 negative). Note that the positive pairs for unbalanced databases are selected randomly from the set of curated positive pairs. Struct2Graph again outperforms its deep-learning counterparts consistently for this unbalanced case. Struct2Graph improvement increases when the ratio between positive and negative pairs becomes increasingly skewed. For instance, when the ratio of positive and negative pairs is 1:10, the precision and recall statistics for the Struct2Graph model are 97.54% and 96.43%, respectively, which are higher by 0.98% and 2.14%, respectively than the performance of the next best deep-learning model, DeepFE-PPI.

Bootstrap resampling yields a very similar conclusion, where the Struct2Graph is again shown to outperform its deep-learning counterparts, as well as the Random Forest classifier, consistently for several unbalanced cases (see Tables 7-10). When the ratio of positive and negative pairs is 1:10, the accuracy, precision and recall statistics for the Struct2Graph model are 99.26%, 97.04% and 95.59%, respectively, which are higher by 0.76%, 0.58% and 7.73%, respectively than the performance of the next best deep-learning model, DeepFE-PPI.

While a ratio of 1 : 10 reflects a significant class imbalance between positive and negative examples, the class imbalance in a protein interactome can potentially be of the order of 1 : 100 or even larger. In the absence of a PPI database (consisting of 3D-structural information) with such huge class imbalance, prevalence-corrected Precision-Recall Curves (PRCs) have been adopted [76] that reduce the false positive rate at the expense of the true positive rate. Figure 3 depicts the prevalence-corrected PRCs on the balanced (1:1) for several PPI classifiers. The computation of precision is suitably modified with *r* = 100 [76] for an expected ratio of 1:100 for positive to negative samples in the real-world data. PRCs best summarize the trade-off between the true positive rate and the positive predictive value (PPV) for a classifier using different probability thresholds. The AUC (area under curve) in Figure 3(i) nearly approaches unity, thus guaranteeing excellent discrimination capability of the proposed Struct2Graph architecture. Figure 4 depicts the prevalence-corrected PRCs for the deep-learning classifiers on the unbalanced (1:10) dataset. As before, the AUC for the Struct2Graph architecture almost approaches unity.

**Figure 3.**
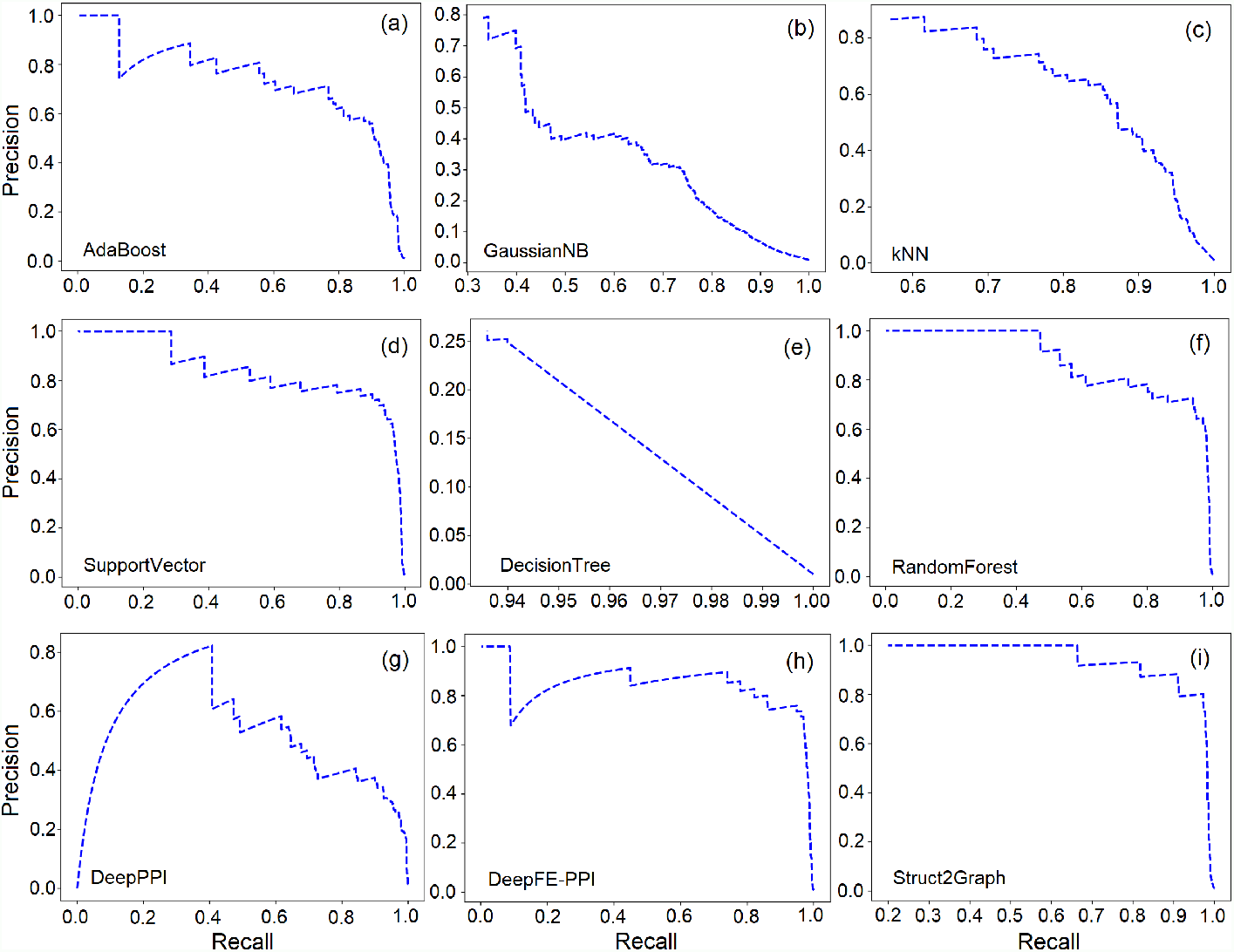
Prevalence corrected precision-recall curves for the balanced database. (a) AdaBoost classifier, (b) GaussianNB classifier, (c) *k*NN classifier, (d) SVC, (e) Decision tree classifier, (f) Random forest classifier, (g) DeepPPI classifier, (h) DeepFE-PPI classifier, (i) Struct2Graph (ours) classifier.

**Figure 4.**
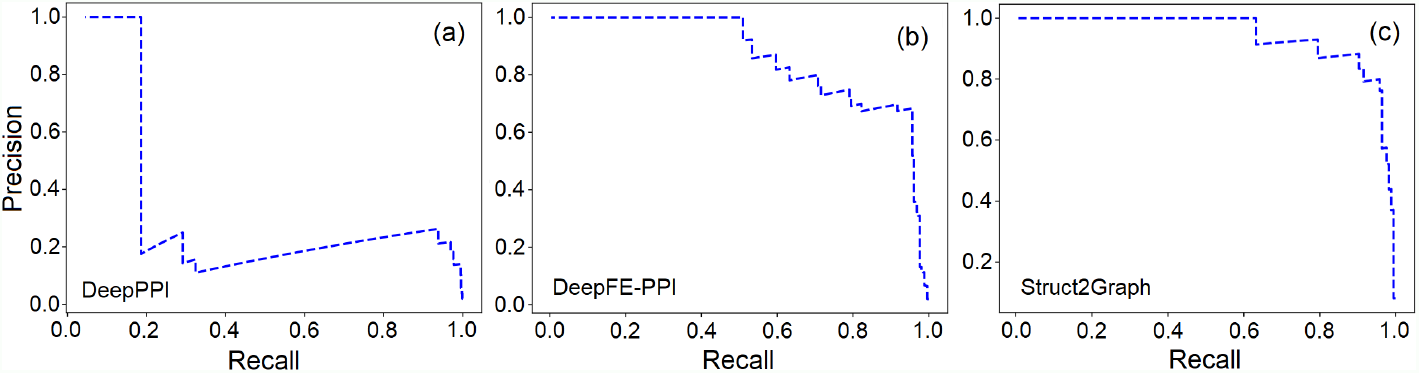
Prevalence corrected precision-recall curves for the unbalanced database. **(a)** DeepPPI classifier, (b) DeepFE-PPI classifier, (c) Struct2Graph (ours) classifier.

### Heterogeneity of the PPI database

The success of any machine learning algorithm is based on the quality of training data it is presented with. Of the 4698 positive and 5036 negative examples spanned across 3677 unique proteins, we first want to make sure that the learning algorithms are not biased towards memorizing the training data [77], since some of the proteins in the database are involved in positive only or negative only interactions. Figure 5 shows the distribution of number of interactions per protein that are involved in positive only interactions. It can be seen that nearly 82% of the unique proteins are involved in four or fewer positive only interactions. Consequently, for a classifier to memorize the training data and not learn to predict positive interactions would be extremely difficult without each protein appearing in a relatively large number of PPI instances.

**Figure 5.**
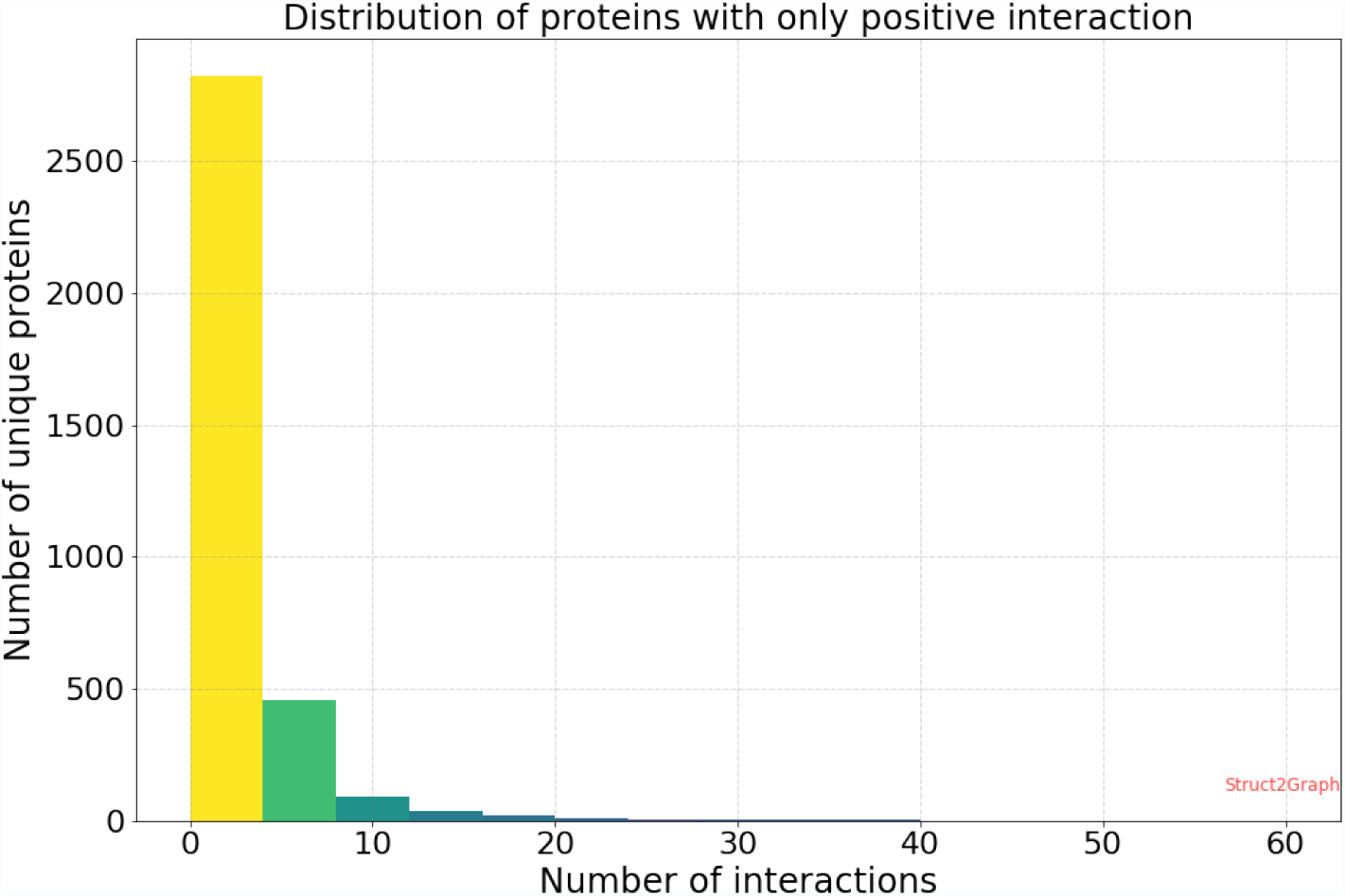
Histogram of proteins with only positive interactions. Of the 3677 unique PDBs, 3453 PDBs are involved in only positive interactions, i.e., among all the protein-protein pair instances in our database, these 3453 proteins do not feature in any non-complex forming instance. Moreover, of the 3453 PDBs with only positive interactions, nearly 82% unique PDBs are involved in fewer than 4 PPI examples. Consequently, for a classifier to memorize data and not “learn” to predict interactions would be extremely difficult without each PDB appearing in a relatively large number of PPI instances.

Similarly, of the 3677 unique proteins, 104 unique proteins are involved in negative only interactions (do not form complexes). Figure 6 shows the distribution of number of interactions per protein that are involved in negative only interactions. As seen in the histogram, the total number of proteins with negative only interactions appearing in more than five PPI examples is very small (81), and comprise of only 2.2% of the entire PDB database considered in our work. Thus, the distribution of data makes it implausible for learning algorithms to perform well just by memorizing the training data. The dataset also consists of 120 unique proteins that are involved in both positive and negative interactions. These 120 unique proteins appear in 6335 PPI instances in our database. Hence, it would be nearly impossible for any classifier to simply memorize the training data, and still be able to predict the interactions almost accurately on the test or validation set.

**Figure 6.**
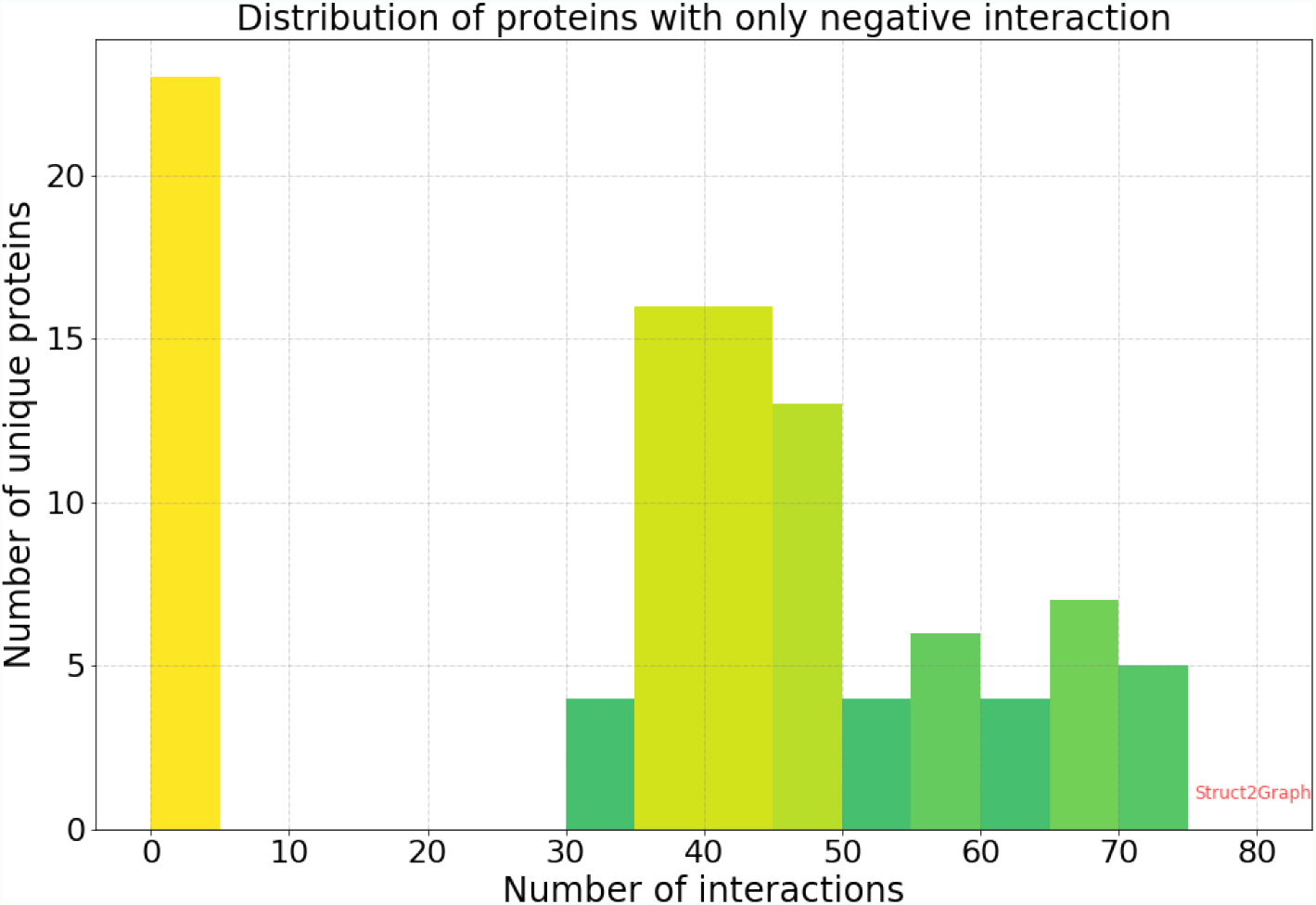
Histogram of proteins with only negative interactions. Of the 3677 unique PDBs, only 104 PDBs are involved in just the negative interactions, i.e., among all the protein-protein pair instances in our database, these 104 proteins do not feature in any complex forming instance. Moreover, of the 104 PDBs, 23 PDBs appear in less than 5 PPI examples. The total number of proteins that are involved in more than 5 PPI examples is a very small number (81), i.e., only 2.2% of the entire PDB database considered in our work.

We further validate this by building a random forest classifier that is supplied with an input vector of length 3677. The specific choice of length of the input vector is directly related to the number of unique proteins in the database. We first create a dictionary of all the unique proteins and note down the order in which these unique proteins appear in the dictionary. Then, each unique protein is represented by a 3677-long unit vector with all but one coordinate being zero. The coordinate corresponding to the order of the protein in the dictionary is marked as 1. For predicting the interaction of two proteins, say proteins A and B, the unit vectors are summed and supplied to the random forest classifier as an input. Recall that the sum operation is permutation invariant, and thus interaction prediction for the pair (protein A, protein B) is identical to that for the pair (protein B, protein A). Table 11 summarizes the performance of the random forest classifiers on balanced and unbalanced datasets, trained using only the labels of protein pairs and disregarding any structural information. In the balanced scenario, the classifier can be trained with a reasonable accuracy of ∼ 91% on both training and test sets. This is still significantly smaller than accuracies obtained using Struct2Graph and other deep-learning based classifiers on the balanced set. However, as the training database is made more realistic (i.e., biased towards significantly abundant negative examples), the performance on the training set drops, while the performance on the test set is completely random (∼ 50%), i.e., the random forest classifier acts like a random predictor. In the extreme scenario, (1:10 ration between positive and negative examples), the training accuracy appears improved, largely because every time the classifier predicts a negative interaction, it is likely to be correct since the training set has an abundance of negative examples. However, on the test set comprising of approximately equal number of positive or negative examples, the prediction accuracy is still around 50% indicating zero learning.

**Table 11.** Memorization test for the PPI database

| Database | Accuracy on Train-Set | Accuracy on Test-Set |
| --- | --- | --- |
| Balanced (1:1) | 91.17% | 91.07% |
| Unbalanced (1:2) | 66.98% | 50.43% |
| Unbalanced (1:3) | 75.25% | 50.23% |
| Unbalanced (1:5) | 83.53% | 50.43% |
| Unbalanced (1:10) | 91.04% | 50.33% |

## Discussion

The success of Struct2Graph is attributed to the thorough analysis of structural 3D information embedded in the form of a graph, which predicts interactions better than sequence-based approaches [13]. In addition, Struct2Graph can potentially identify residues that likely contribute to the formation of the protein-protein complex. This is achieved by considering the probability tuples *{*(*p*_*i*_, *p*_*j*_)*}* of different amino acids during the knowledge selection process described in Equation (3). These probabilities capture the relative importance of amino acids and thus reflect different amino acids’ contributions towards interaction prediction. The amino acids with large relative probabilities (top 20%) are identified as important for the formation of the protein-protein complex. This importance can stem from either direct participation in the interaction process (i.e., binding site) or indirectly through contribution to appropriate protein folding that allows formation of the correct binding site geometry.

A demonstration of the potential of Struct2Graph to identify specific interaction sites was performed on two example cases (neither part of the training set) with well-described interacting residues from protein pairs in the literature. Specifically, we studied two different interaction types: 1) A protein with multiple ligands competing for the same binding area [78]; and 2) A dynamic protein-protein adhesion interaction [79]. The reported interacting residues in these complexes are compared with the Struct2Graph’s highest probability residues (top 20%) using standard 2×2 confusion matrices. In aggregate (i.e., two case examples with a total of three interacting pairs), Struct2Graph identifies interacting residues with 30% sensitivity, 89% specificity, and 87% accuracy. It should be noted that these protein pair examples are not in the training set, and Struct2Graph identifies these residues through its knowledge selection process in a completely unsupervised manner. Besides, as noted above, the identified residues could be critical for ensuring correct protein folding conformation and therefore *indirectly* important for predicting binding, but not captured by traditional analysis that focuses only on the specific interacting residues identified in the literature. Detailed results for each example are described:

### 1) HMGB1 and PSM*α*_1_ compete for binding TLR4

Phenol soluble modulins (PSMs), short, amphipathic, helical peptides [80], play a crucial role in *Staphylococcus aureus* virulence, one of the most common causes of human bacterial infections worldwide [81]. *S. aureus* has seven PSMs (PSM*α*_1_ −*α*_4_, PSM*β*_1_ −*β*_2_, and *δ*-toxin) which have multiple functions including, cytolysis, biofilm structuring, and inflammatory activation via cytokine release and chemotaxis. PSMs specifically trigger the release of high mobility group box-1 protein (HMGB1). Toll-like receptor-4 (TLR4) interacts with HMGB1 activating nuclear factor NF-κB and proinflammatory cytokines production [82]. However, *S. aureus* PSMs*α*_1_ *− α*_3_ significantly inhibit HMGB1-mediated phosphorylation of NF-κB by competing with HMGB1 via interactions with the same residues of TLR4 domain [78]. As such, the specific interacting residues for these pairs HMGB1:TLR4 (2LY4 : 3FXI) and PSM*α*_1_:TLR4 (5KHB : 3FXI) have been well described [78].

Struct2Graph identifies interacting residues of the HMGB1:TLR4 pair with 90% accuracy in which the top 9 predicted residues for TLR4 fall within the reported active cavity (residues rank 336-477). In addition, among the top 20% predicted residues of HMGB1 were the specific interacting residues Tyr^16^ and Lys^68^. For the PSM*α*_1_:TLR4 pair, Struct2Graph identifies interacting residues with 92% accuracy. Again the top predicted residues fall within the previously identified TLR4 active cavity (336-477). For PSM*α*_1_, interacting residues Gly^2^ and Val^10^ were correctly identified. While the overall sensitivity for detecting an interacting residue is ∼ 20% for this example, Struct2Graph was able to predict that PSM*α*_1_ interacts with TLR4 in the same area as the HMGB1 binding site. More specifically, the predicted binding sites for both on TLR4 have 94% concordance. Figures 7a and 7b shows the residues predicted to be essential and highlights how Struct2Graph predicts a similar site for both interactions.

**Figure 7.**
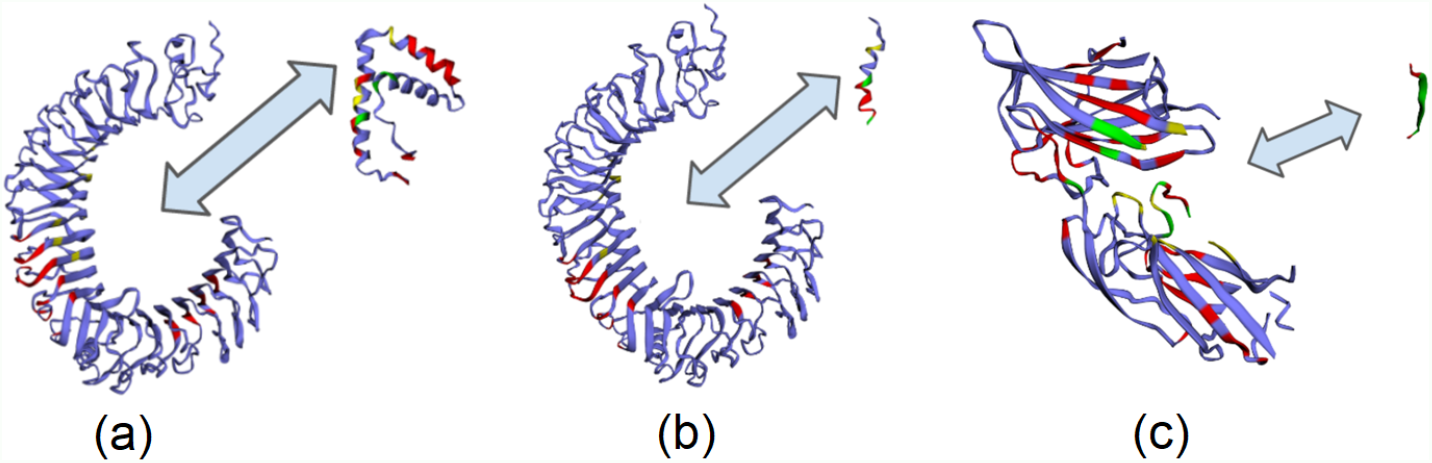
Important residue prediction by Struct2Graph for three example scenarios. (a) TLR4 with HMGB1, (b) TLR4 with PSM*α*_1_, (c) SdrG and Fibrinogen adhesion. The different colored residues encode different information: (i) Red: Top-20% residues identified important by Struct2Graph, (ii) Yellow: Actual binding site not identified to be important by Struct2Graph, (iii) Green: True binding site overlapping with a residue identified important by Struct2Graph, (iV) Purple: neither important, nor actual interaction site. Recall that both HMGB1 and PSM*α*_1_ are known to compete for the same binding sites on TLR4, and this gets reflected in the Struct2Graph predictive analysis as well.

### 2) SdrG-Fibrinogen Adhesion

Microbial attachment to host tissues is a crucial step in most bacterial infections. Gram-positive pathogens such as Staphylococci, Streptococci, and Enterococci contain multiple cell wall-anchored proteins that act as an adhesin to mediate bacterial attachment to host tissues. These adhesin mediating interactions have been termed MSCRAMMs (microbial surface components recognizing adhesive matrix molecules) [83]. SdrG is an MSCRAMM of *Staphylococcus epidermidis* that binds to the *Bβ* chain of human fibrinogen (Fg) via dynamic “dock, lock, and latch” mechanism [79].

Struct2Graph was used to evaluate the interaction between SdrG (PDB:r19A) and a synthetic peptide with homologous sequence to its binding site in Fg (PDB:r17C). Interacting residues between SdrG and the synthetic Fg peptide homolog were predicted with 75% accuracy. Among the high probability residues identified in SdrG were 9 exact matches to those in the literature [79]. This included, Pro^337^, Ser^338^, Leu^340^, Phe^344^, Gln^425^, Ser^437^, Tyr^577^, Asp^578^, and Asn^579^. Figure 7c shows the residues predicted to be essential for the interaction.

These results show that Struct2Graph provides insight into key residues involved in the protein-protein interaction without any training data on the specific nature of these interactions. A complete summary of the residues identified by Struct2Graph for the preceding examples is included in the supplementary material (see Additional File 1). Any high probability residues identified but not confirmed as directly interacting may have indirect effects through maintaining appropriate 3D conformation of the protein.

In addition to these specific binding examples, we consider the ability of our attention mechanism to predict useful residues across a broader dataset. Our attention mechanism does not necessarily predict interaction sites, but rather residues which are important to protein interactions regardless of their proximity to the interface. It has been observed that residues throughout the entire peptide chain can drive interactions [84]. Therefore, the attention mechanism will identify residues regardless of location that significantly alter interaction propensity. To demonstrate this, we analyze the single amino acid variation (SAV) dataset presented in [85] (please refer to the supporting information `Additional File` 2 for the human SAV dataset). The authors in [85] performed a large-scale structural analysis of human single amino acid variations (SAVs) and demonstrated that disease-causing mutations are preferentially located within the interface core, as opposed to the rim. Their work analyzed a total of 3282 disease-causing SAVs and 1699 benign polymorphisms occurring in 705 proteins. It is established that the disease-causing SAVs were 49% more likely to occur in the interface core rather than the rim and were 72% more likely to occur in the interface core than in the non-interacting protein surface, thus clearly demonstrating a different contribution of core and rim regions to human disease. On the other hand, 78.7% of polymorphisms were found to reside within surface-accessible residues (241 in interface residues and 1096 in surface non-interface residues), i.e, polymorphisms are less likely to be located in the interface core compared to the rim.

Since the work in [85] primarily dealt with human database, there is sufficient overlap between their dataset and the PPI database used in our manuscript. Of the overlapping 2724 disease-causing SAVs (spanning across 342 unique proteins) and 1364 polymorphisms (spanning across 528 unique proteins), our attention mechanism identifies 33.55% of all disease-causing SAVs as important (attention weights within top-20%), while 85.30% of all polymorphisms are identified as *un*important by the proposed attention mechanism, indicating significant overlap between the previously established SAV study and the important residues identified by the proposed attention mechanism.

## Conclusion

Struct2Graph, a GCN-based mutual attention classifier, to accurately predict interactions between query proteins exclusively from 3D structural data is proposed. Since the prior study showed that the geometrical and graph theoretical descriptors may be sufficient for description of PPI [13], Struct2Graph does not directly use descriptors, such as sequence information, hydrophobicity, surface charge and solvent accessible surface area, and thus can be generalized to a broader class of nanoscale structures that can be represented in similar fashion. This study demonstrates that a relatively low-dimensional feature embedding learned from graph structures of individual proteins outperforms other modern machine learning classifiers based on global protein features. Our GCN-based classifier achieves state-of-the-art performance on both balanced and unbalanced datasets.

Moreover, the mutual attention mechanism provides insights into important residues that are likely to contribute towards interaction through direct or indirect participation. This is achieved through its knowledge selection process in a completely unsupervised manner. The identification of important residues is tested for two different interaction types: (a) Protein with multiple ligands competing for the same binding area, (b) Dynamic protein-protein adhesion interaction. Struct2Graph identifies interacting residues with 30% sensitivity, 89% specificity, and 87% accuracy. Finally, through the analysis of single amino acid variations, the attention mechanism shows preference for disease-causing residue variations over benign ones, demonstrating that it is not limited to interface residues. This connection between the unsupervised discovery of interaction sites and graph representation of proteins is possible thanks to the somewhat limited type of atoms and bond patterns that commonly occur in such molecules, which makes it possible to characterize properties on local atomistic arrangements. Overall, the proposed framework is general and, while subject to availability of corresponding training data, can be made to predict other kinds of complex sets of collective supramolecular interactions between proteins and nanoscale species of different chemical composition.

## Funding and Acknowledgements

AV, JSV, PE, MB, AM and SK acknowledge the support from the BlueSky Initiative from the University of Michigan College of Engineering. AOH acknowledges the support from ARO W911NF-19-1-0269 and ARO W911NF-14-1-0359. AV and JS acknowledge the support from DARPA HR00111720067. NAK expresses gratitude to Vannewar Bush DoD Fellowship ONR N000141812876. The funders had no role in study design, data collection and analysis, decision to publish, or preparation of the manuscript.

## Abbreviations

PPI: Protein-Protein Interaction
GAT: Graph Attention Network
GCN: Graph Convolutional Network

## Availability of data and materials

The source code, along with the PPI database, is available at https://github.com/baranwa2/Struct2Graph.

## Authors’ contributions

**Mayank Baranwal:** Methodology, Software, Analysis, Writing - original draft. **Abram Magner:** Methodology, Software, Analysis. **Jacob Saldinger:** Validation of PPI results, Comparison with existing methods. **Emine S. Turali-Emre:** Data curation, Validation, Writing - PPI database and interaction site prediction sections. **Shivani Kozarekar:** Data curation. **Paolo Elvati:** Methodology - representation and properties of molecules, Conceptualization, Writing - review & editing. **J. Scott VanEpps:** Writing - review & editing, Supervision - PPI database. **Nicholas A. Kotov:** Methodology - graph representation of proteins and other nanostructures, Writing - review & editing, Supervision - PPI database. **Angela Violi:** Conceptualization, Writing - review & editing, Supervision. **Alfred O. Hero:** Conceptualization, Writing - review & editing, Supervision.

## Additional Files

Additional File 1 — Supplementary Material

This file contains details of important residues identified by Struct2Graph for test pairs discussed in Figure 7. Additionally, the supporting information also sheds light on creating protein embeddings using Struct2Graph.

## Footnotes

[1] Based on the pairwise homology analysis comprising of 3677 unique proteins in our database, only 0.3% of the proteins were found to have BLAST e-value *<* 0.05 and 0.26% has *<* 0.001, indicating statistically insignificant homologous relationships.

