## Additional File 1 for "Struct2Graph: A graph attention network for structure based predictions of protein-protein interactions"

### Supplementary Material

---

#### Details Of Important Residues

A demonstration of the potential of Struct2Graph to identify specific interaction sites was performed on two example cases (neither part of the training set) with well-described interacting residues from protein pairs in the literature. Specifically, we studied two different interaction types: 1) A protein with multiple ligands competing for the same binding area [1]; and 2) A dynamic protein-protein adhesion interaction [4]. The reported interacting residues in these complexes are compared with the Struct2Graph's highest probability residues (top 20%) using standard 2x2 confusion matrices. In aggregate (i.e., two case examples with a total of three interacting pairs), Struct2Graph identifies interacting residues with 30% sensitivity, 89% specificity, and 87% accuracy. It should be noted that these protein pair examples are not in the training set, and Struct2Graph identifies these residues through its knowledge selection process in a completely unsupervised manner. Besides, as noted above, the identified residues could be critical for ensuring correct protein folding conformation and therefore *indirectly* important for predicting binding, but not captured by traditional analysis that focuses only on the specific interacting residues identified in the literature. Detailed results for each example are described:

1) **HMGB1 and PSM$\alpha_1$ compete for binding TLR4**: Phenol soluble modulins (PSMs), short, amphipathic, helical peptides [2], play a crucial role in *Staphylococcus aureus* virulence, one of the most common causes of human bacterial infections worldwide [5]. *S. aureus* has seven PSMs (PSM$\alpha_1-\alpha_4$, PSM$\beta_1-\beta_2$, and $\delta$-toxin) which have multiple functions including, cytolysis, biofilm structuring, and inflammatory activation via cytokine release and chemotaxis. PSMs specifically trigger the release of high mobility group box-1 protein (HMGB1). Toll-like receptor-4 (TLR4) interacts with HMGB1 activating nuclear factor NF-κB and proinflammatory cytokines production [6]. However, *S. aureus* PSMs$\alpha_1-\alpha_3$ significantly inhibit HMGB1-mediated phosphorylation of NF-κB by competing with HMGB1 via interactions with the same residues of TLR4 domain [1]. As such, the specific interacting residues for these pairs HMGB1:TLR4 (2LY4 : 3FXI) and PSM$\alpha_1$:TLR4 (5KHB : 3FXI) have been well described [1].

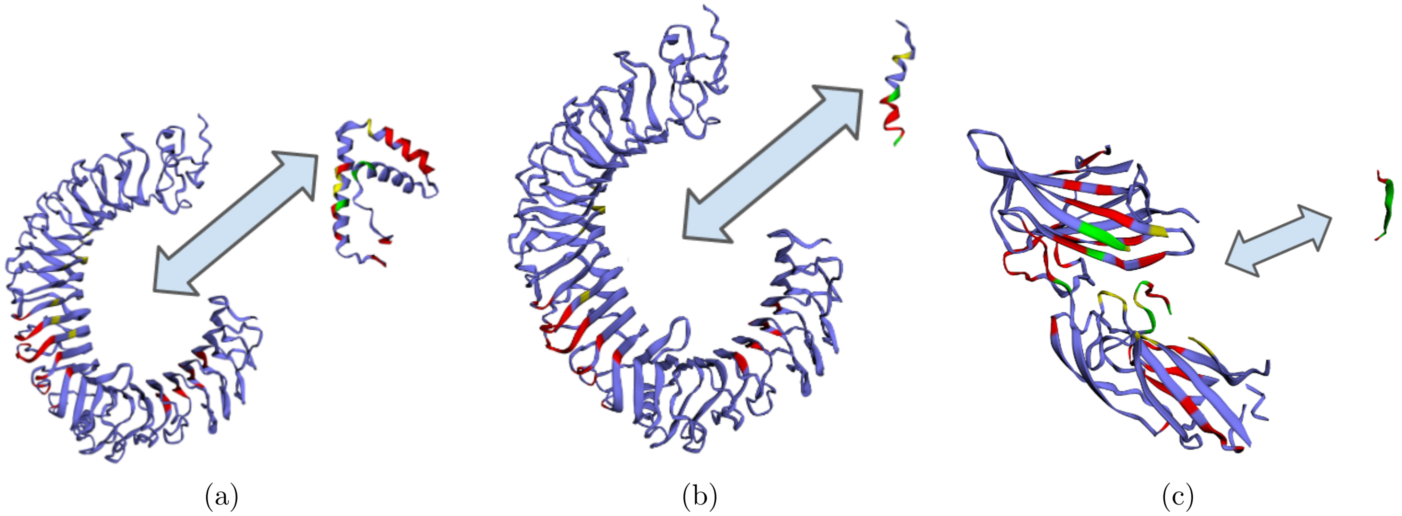


Figure 1: **Important residue prediction by Struct2Graph for three example scenarios.** (a) TLR4 with HMGB1, (b) TLR4 with PSM$\alpha$, (c) SdrG and Fibrinogen adhesion. The different colored residues encode different information: (i) Red: Top-20% residues identified important by Struct2Graph, (ii) Yellow: Actual binding site not identified to be important by Struct2Graph, (iii) Green: True binding site overlapping with a residue identified important by Struct2Graph, (iV) Purple: neither important, nor actual interaction site. Recall that both HMGB1 and PSM$\alpha_1$ are known to compete for the same binding sites on TLR4, and this gets reflected in the Struct2Graph predictive analysis as well.

### Producing Mol2Vec-Like Embeddings Using Struct2Graph

Struct2Graph can be used to produce protein embeddings, similar to molecular embeddings produced by Mol2Vec. The embeddings are simply the output of the last layer of the graph convolutional layer averaged across the size of the graph. In the source code provided on Github (`https://github.com/baranwa2/Struct2Graph/blob/master/k-fold-CV.py`), the outputs $xs1$ and $xs2$ from the 'gnn' are averaged along the first-dimension to produce Mol2Vec-like embeddings.
